# Development of Microstructural and Morphological Cortical Profiles in the Neonatal Brain

**DOI:** 10.1101/2020.01.14.906206

**Authors:** Daphna Fenchel, Ralica Dimitrova, Jakob Seidlitz, Emma C. Robinson, Dafnis Batalle, Jana Hutter, Daan Christiaens, Maximilian Pietsch, Jakki Brandon, Emer J. Hughes, Joanna Allsop, Camilla O’Keeffe, Anthony N. Price, Lucilio Cordero-Grande, Andreas Schuh, Antonios Makropoulos, Jonathan Passerat-Palmbach, Jelena Bozek, Daniel Rueckert, Jo V. Hajnal, Armin Raznahan, Grainne McAlonan, A. David Edwards, Jonathan O’Muircheartaigh

## Abstract

In the perinatal brain, regional cortical architecture and connectivity lay the foundations for functional circuits and emerging behaviour. Interruptions or atypical development during or before this period may therefore have long-lasting consequences. However, to be able to investigate these deviations, we need a measure of how this architecture evolves in the typically developing brain. To this end, in a large cohort of 241 term-born infants we used Magnetic Resonance Imaging to estimate cortical profiles based on morphometry and microstructure over the perinatal period (37-44 weeks post-menstrual age, PMA). Using the covariance of these profiles as a measure of inter-areal network similarity (Morphometric Similarity Networks; MSN), we clustered these networks into distinct modules. The resulting modules were consistent and symmetric, and corresponded to known functional distinctions, including sensory-motor, limbic and association regions and were spatially mapped onto known cytoarchitectonic tissue classes. Posterior (parietal, occipital) regions became more morphometrically similar with increasing PMA, while peri-cingulate and medial temporal regions became more dissimilar. Network strength was associated with PMA: Within-network similarity increased over PMA suggesting emerging network distinction. These changes in cortical network architecture over an eight-week period are consistent with, and likely underpin, the highly dynamic behavioural and cognitive development occurring during this critical period. The resulting cortical profiles might provide normative reference to investigate atypical early brain development.

## Introduction

Brain maturation over the perinatal period is rapid and complex. Although the majority of neurons are at their terminal location, synaptogenesis and myelination are ongoing, and (limited) migration of interneurons continues (Paredes et al., 2016; Petanjek et al., 2011). In the cortex, these processes are coordinated with sensory cortex and pathways developing earliest, prefrontal and association areas later (Flechsig, 1901; Huttenlocher, 1979). This developing architecture and connectivity is critical for efficient functional signalling, supporting and enabling the later development of cognitive and behavioural abilities.

Although alterations in perinatal neurodevelopmental processes have been associated with later cognitive and behavioural difficulties, quantifying myelo- or cytoarchitecture anatomical circuit development in the living human neonate is challenging. A relatively simple hypothesis of cortical connectivity is “similar prefers similar” (Goulas et al., 2016), that is areas with similar cytoarchitecture preferentially connect. In this context, Magnetic Resonance Imaging (MRI)-based methods such as regional structural covariance (Evans, 2013) offer a proxy measure of brain connectivity, with structural similarity of spatially distinct regions of cortex reflecting coordinated maturation (Alexander-Bloch, Giedd & Bullmore, 2013). This inter-regional similarity is supported by similar genetic or maturational profiles (Alexander-Bloch et al. 2013), transcriptomic profiles (Yee et al., 2018), or associated with changes in disease (e.g., Zuo et al., 2018). Together with complementary information from MRI estimates of structural (tractography) and functional connectivity (Gong et al., 2013), regional similarity could capture more accurately the brain’s structural and functional organization.

Understanding early structural regional similarity can shed light on how the brain develops into an efficient multi-functional system (Cao et al., 2016) and allow the detection of perturbations in normal development (Di Martino et al., 2014; Morgan et al., 2018). Structural Covariance Networks (SCNs) based on measures of grey matter (GM) volume, cortical thickness (CT), cortical folding, and fiber density in the first two years of life have been described (Fan et al., 2011; Geng et al., 2017; Nie et al., 2014). As different anatomical measures are capturing distinct developmental processes (Rakic, 1988) and are controlled by different genetic mechanisms (Chen et al., 2013; Panizzon et al., 2009), different measures of cortical growth and development show specific spatial and temporal patterns in early childhood (Gilmore et al. 2012; Li et al. 2013; Lyall et al., 2015; Nie et al., 2014). Therefore, as might be expected, the resulting single-feature SCNs can be inconsistent. Because the possibility to generate individual SCNs is limited (Tijms et al., 2012), SCNs must be defined at the group level and they cannot provide person-level information on anatomical relatedness between brain regions.

An alternative approach using multiple anatomical measures to elucidate regional structural similarity has recently been examined in adolescents and adults (Li et al., 2017; Seidlitz et al., 2018). Compared to SCNs derived from a single technique, this approach allows for the construction of networks for individual subjects, rather than over a group, enabling the assessment of individual variability masked by group templates or case-control studies (Seghier & Price, 2018). The resulting Morphometric Similarity Networks (MSNs) have superior spatial consistency with cortical cytoarchitecture compared to structural networks based on diffusion imaging or on one structural measure (CT) (Seidlitz et al., 2018). Furthermore, cortical regions shown to be connected (i.e., similar) based on this approach have complementary expression of human supra-granular enriched genes (HSE) (Romero-Garcia et al., 2018; Seidlitz et al., 2018). In adults, these networks are also sensitive to alternations in common (Morgan et al., 2019) and rare (Seidlitz et al., 2019) neurodevelopmental disorders. They also have neurobiological specificity; cortical MSNs differ between patients with genetic syndromes and these changes are closely aligned with the regional expression of disorder-related genes (Seidlitz et al., 2019). Recent work in neonates using a variant of MSNs have successfully predicted postmenstrual age (PMA) at scan and differentiated infants born prematurely, with superior performance compared to using single predictive measures (Galdi et al., 2019).

The aim of this work therefore was to examine the multi-morphometric cortical organization of the newborn brain across the perinatal period from 37 to 44 weeks PMA in a large sample of 241 infants. Using cortical MSNs constructed from both structural / morphological and microstructural diffusion parameters, we explored brain development over this important period of rapid growth. We used data-driven clustering approaches to describe the modular structure of the brain, demonstrating a mapping to known functional and cytologic regions and significant changes with increasing PMA.

## Methods

### Subjects

This work included a sample of neonates participating in the Developing Human Connectome Project (dHCP); (http://www.developingconnectome.org/), scanned at the Newborn Imaging Centre at Evelina London Children’s Hospital, London, UK. Images are available for download and analysis at the project website. This project has received ethical approval (14/LO/1169, IRAS 138070), and written informed consent was obtained from parents. Two-hundred and forty-one singleton normal term-born subjects (born at between 37 and 42 weeks; 128 males) with good quality structural and diffusion MR images acquired at PMA 40.92±1.58 weeks (mean±sd), range 37.43-44.71 weeks) were included in this analysis. No major brain abnormalities were detected in review of the MRI data by a neonatal neuroradiologist.

### Image Acquisition

MR images were acquired on a 3T Phillips Achieva scanner without sedation, using a dedicated 32-channel neonatal head coil system (Hughes et al., 2017). Acquisition and reconstruction followed optimized protocols for structural images (Cordero-Grande et al., 2018; Cordero-Grande et al., 2016; Kuklisova-Murgasova et al., 2012), and multi-shell High Angular Resolution Diffusion Imaging (HARDI) (Hutter et al., 2018; Tournier et al., 2015). T2-weighted (T2w) images were obtained using a Turbo Spin Echo (TSE) sequence, acquired in two stacks of 2D slices (in sagittal and axial planes), using parameters: TR=12s, TE=156ms, SENSE factor 2.11 (axial) and 2.58 (sagittal) with overlapping slices (resolution 0.8×0.8×1.6mm^3^). T1-weighted (T1w) images were acquired using an IR (Inversion Recovery) TSE sequence with the same resolution using TR=4.8s, TE=8.7ms, SENSE factor 2.26 (axial) and 2.66 (sagittal). Structural images were reconstructed to a final resolution 0.5×0.5×0.5mm^3^ using slice-to-volume registration. Diffusion images were obtained using parameters TR=3800ms, TE=90ms, SENSE factor=1.2, multiband factor=4, resolution 1.5×1.5×3.0mm^3^ with 1.5mm slice overlap. Diffusion gradient encoding included images collected at b=0s/mm^2^ (20 repeats), b=400s/mm^2^ (64 directions), b=1000s/mm^2^ (88 directions), b=2600s/mm^2^ (128 directions), and images were reconstructed to a final resolution of 1.5×1.5×1.5mm^3^.

### Image Processing

Cortical surface processing and extraction of individual cortical features followed the pipeline described in (Makropoulos et al., 2018). Briefly, motion- and bias-corrected T2w images were brain extracted and segmented. White, pial and midthickness surfaces were extracted, inflated and projected onto a sphere. This was followed by estimation of cortical features including CT, pial surface area (SA), mean curvature (MC), and a gross proxy of myelin content, defined as the ratio between the T1w/T2w images (Glasser & Van Essen, 2011), performed after registering the individual T1 and T2 images together, henceforth “myelin index (MI)”. All brains were aligned to the 40-week dHCP surface template (Bozek et al., 2018) using Multimodal Surface Matching (MSM) (Robinson et al., 2013, 2014), run with higher order regularization constraints (Robinson et al., 2018), to match coarse scale cortical folding (sulcal depth) maps. All other metrics were resampled to the template using this transformation and adaptive barycentric resampling (implemented using Human Connectome Project (HCP) tools, Connectome Workbench (https://www.humanconnectome.org/software/connectome-workbench).

Diffusion images were denoised (Veraart et al. 2016);, Gibbs-ringing suppressed (Kellner et al., 2016);, and corrected for subject motion- and image distortion with slice-to-volume reconstruction in the multi-shell spherical harmonics and radial decomposition (SHARD) basis (Christiaens et al., 2019a), as described in (Christiaens et al. 2019b) and using an image-based field map (Andersson, Skare & Ashburner, 2003). A tensor model for diffusion data was fitted using a single shell (b=1000), and fractional anisotropy (FA) and mean diffusivity (MD) maps were generated using MRtrix3 (Tournier et al., 2019). Neurite density index (NDI) and orientation dispersion index (ODI) maps were calculated using the NODDI toolbox (Zhang et al., 2012) as previously applied in the neonatal brain (Batalle et al., 2019). Diffusion maps were registered onto individual T2w images using FSL’s epi_reg (FLIRT) (https://fsl.fmrib.ox.ac.uk), and then projected onto cortical surface using Connectome Workbench in order to sample imaging features with the same spatial representation. All raw images were visually inspected for motion or image artefact and artefacted data excluded, and processed images were inspected for registration errors.

### MSNs construction

Cortical regions were defined using approximately equal-sized cortical parcellations with Voronoi decomposition at different granularities (n=50,100,150,200,250,300). This option was chosen over parcellation with a pre-existing atlas in order to minimize the effect of variable regional size when calculating the MSNs. Network construction followed steps introduced before (Li et al., 2017; Seidlitz et al., 2018): for each subject, an eight-feature vector including mean normalized (to account for feature variation) values of CT, MC, MI, SA, FA, MD, NDI and ODI characterizing each node. Correlation (Pearson’s *r*) between the eight-feature vector for every two pairs of regions was calculated using Matlab, resulting in an *n regions × n regions* similarity matrix for each subject. A group structural similarity matrix was produced by averaging the 241 individual similarity matrices (Figure 1) and clustering was performed with affinity propagation. This process was also applied for a ‘leave-one-out’ analysis, where similarity was based on a series of seven-feature vectors, leaving one feature out each time. In addition, typical SCNs for the individual features which are much more common in the literature, were also generated by correlating each pair of regions across all subjects, resulting in a group similarity matrix that is based on a single morphometric feature. The values used in this analysis were the raw, un-normalized values.

**Figure 1.**
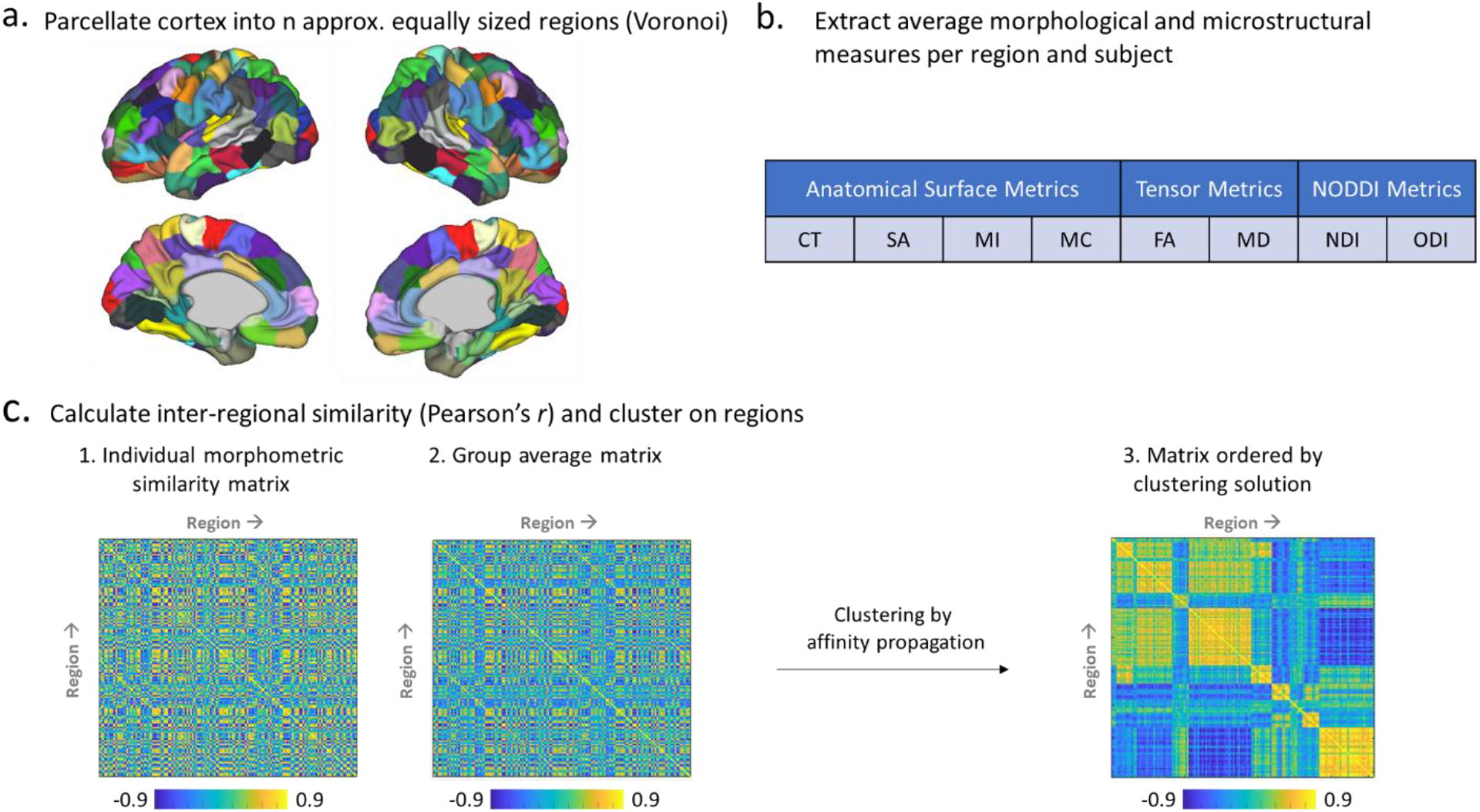
Pipeline for clustering of MSNs in the neonatal brain: a. Cortical regions are defined using Voronoi tessellation of the cortical surface; b. Each individual region is characterized by an eight-feature vector including averaged normalized values of cortical thickness (CT), mean curvature (MC), myelin index (MI), surface area (SA), fractional anisotropy (FA), mean diffusivity (MD), neurite density index (NDI) and orientation dispersion index (ODI); c. Each pair of regions is correlated using Pearson’s *r*, resulting in a subject specific similarity matrix with size n regions × n regions, that are then averaged to create a group mean similarity matrix and the resulting group matrix is clustered using affinity propagation algorithm to examine network modularity.

### Association between MSNs with PMA at scan and sex

To illustrate the effect of PMA on the individual morphometric features, Spearman’s correlation (Spearman’s rho, *ρ*) between the averaged regional single features and PMA was calculated. The effects of PMA on inter-nodal similarity was examined by non-parametric correlation between PMA and the inter-nodal edge-strength. In addition we investigated the same correlation with *mean* node strength (the *average* of a node’s edge-strength with every other node). Sex differences in MSNs for individual edge-strength and mean nodal edge-strength were explored using the Mann-Whitney test.

### MSNs clustering analysis

To investigate network modularity, the group similarity matrix was clustered using affinity propagation (Frey & Dueck, 2007 https://psi.toronto.edu/?q=tools). This clustering method has two merits in this current work: First, clustering could be performed without thresholding the group matrix, therefore avoiding information loss due to binarization, and second, it returns the optimal number of clusters (k) for a given preference value. In our case we used the median of the similarity matrix as preference value, making no prior assumptions, as suggested by Frey & Dueck, and this allowed the exploration of node density on the number of modules. In addition, we also performed the same clustering approach but restricted the solution to a k=7, one less than the number of features from which the MSN was derived.

To measure robustness of the MSN clusters to the type of MRI modalities included, and to rank the importance of input modalities to the eventual solution, agreement between the clustering solution for the eight-feature regional vector and the clustering solutions for seven-feature regional vector (‘leave-one-out analysis’) and clustering solutions for the eight single features (i.e., SCNs) was examined using normalized variation of information (Meilă, 2007; Reichart & Rappoport, 2009) using the ‘nvi’ code from the Pattern Recognition and Machine Learning Toolbox for Matlab (http://prml.github.io/). As in the leave-one-feature out analysis, we fixed k=7.

As MSNs in adults have been reported to correspond with cortical cytoarchitecture (Seidlitz et al., 2018), we sought to examine the overlap between the resulting clusters with gross von Economo cytoarchitectural classifications (seven classes) (von Economo, 1925) as in Whitaker et al. (2016), using the Dice coefficient to estimate overlap.

#### Association between MSN clusters and PMA at scan

To investigate developing network integration and segregation, for each cluster the averaged edge-strength of connections within a cluster (with higher values indicating cluster distinction within the entire network), and the averaged edge-strength of connections between nodes within the cluster and nodes external to that cluster (indicating integration or segregation of that cluster) were calculated. Spearman’s *ρ* coefficients were calculated for changes in these modular integration and segregation measures against PMA.

## Results

### PMA at scan and sex effects

#### PMA-related associations with single-feature maps

The correlation between single-feature maps and PMA at scan revealed metric-specific PMA associations: SA and MI had a brain-wide positive association with PMA, followed by CT, ODI and NDI that also showed a positive association, but to a lesser extent. FA exhibited both positive and negative associations with PMA, while MD displayed only a negative association. Significant correlations with MC results were sparser, seen in limited perisylvian, frontal and temporal regions (Figure 2).

**Figure 2.**
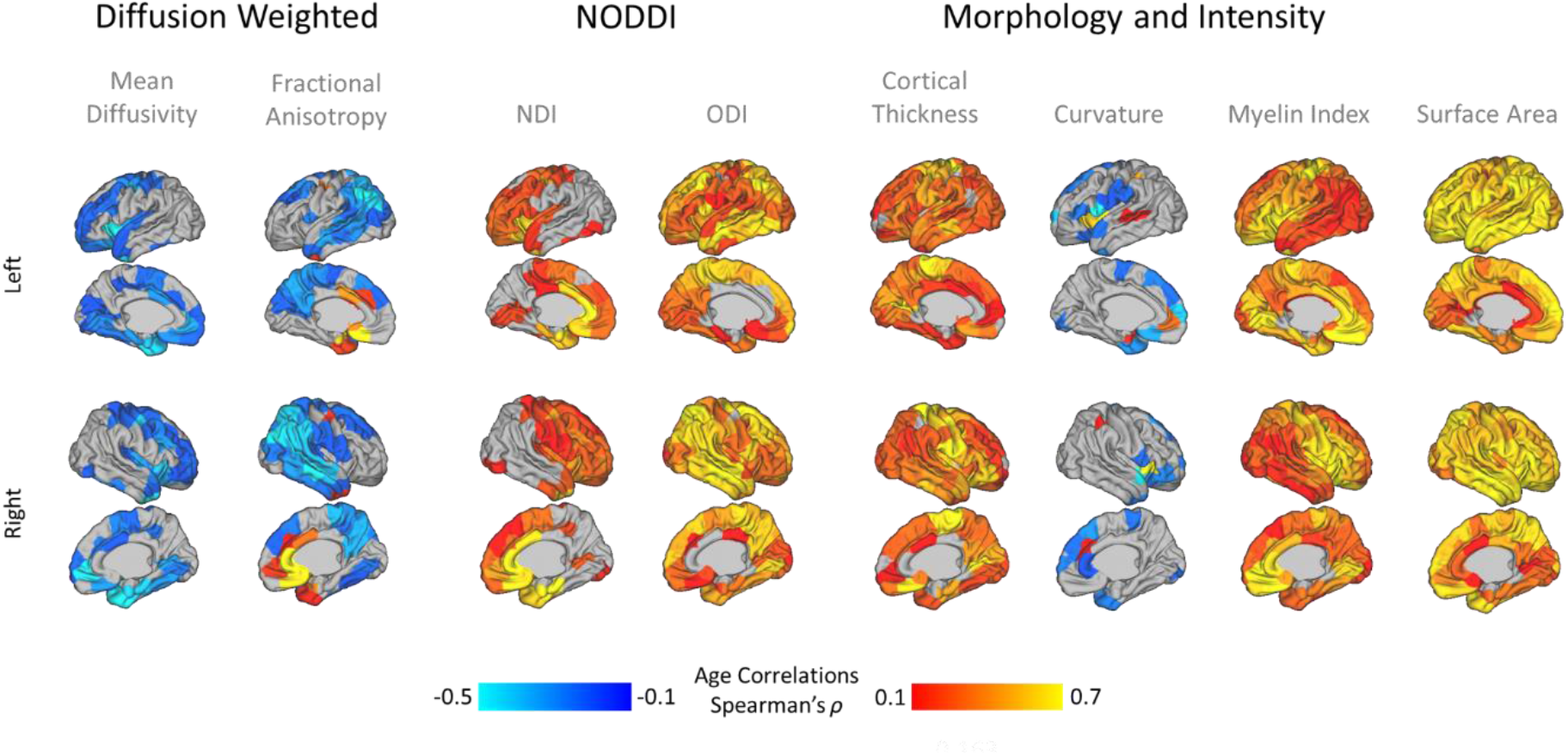
Association between single-feature maps and postmenstrual age (PMA) at scan (results shown corrected for false discovery rate (FDR) at 5%). Positive correlations are marked in red, and negative correlations in blue. NDI-neurite density index, ODI-orientation dispersion index.

#### PMA- and sex-related associations with MSNs

The correlation between inter-regional edge-strength of MSNs and PMA at scan showed both positive and negative associations throughout the brain. They numbered less in frontal and anterior temporal regions (see Figure 3a for a node by node count of significant edges). For mean nodal edge-strength, this spatial gradient was clearer, with anterior cingulate and limbic regions negatively associated with PMA and lateral and medial parietal positively associated (Figure 3b). Individual edge-strength did show some sex associations (Figure 3c), though less extensive, with higher edge-strength in lateral frontal and temporal regions in males and cingulum and medial temporal regions in females (Figure 3d).

**Figure 3.**
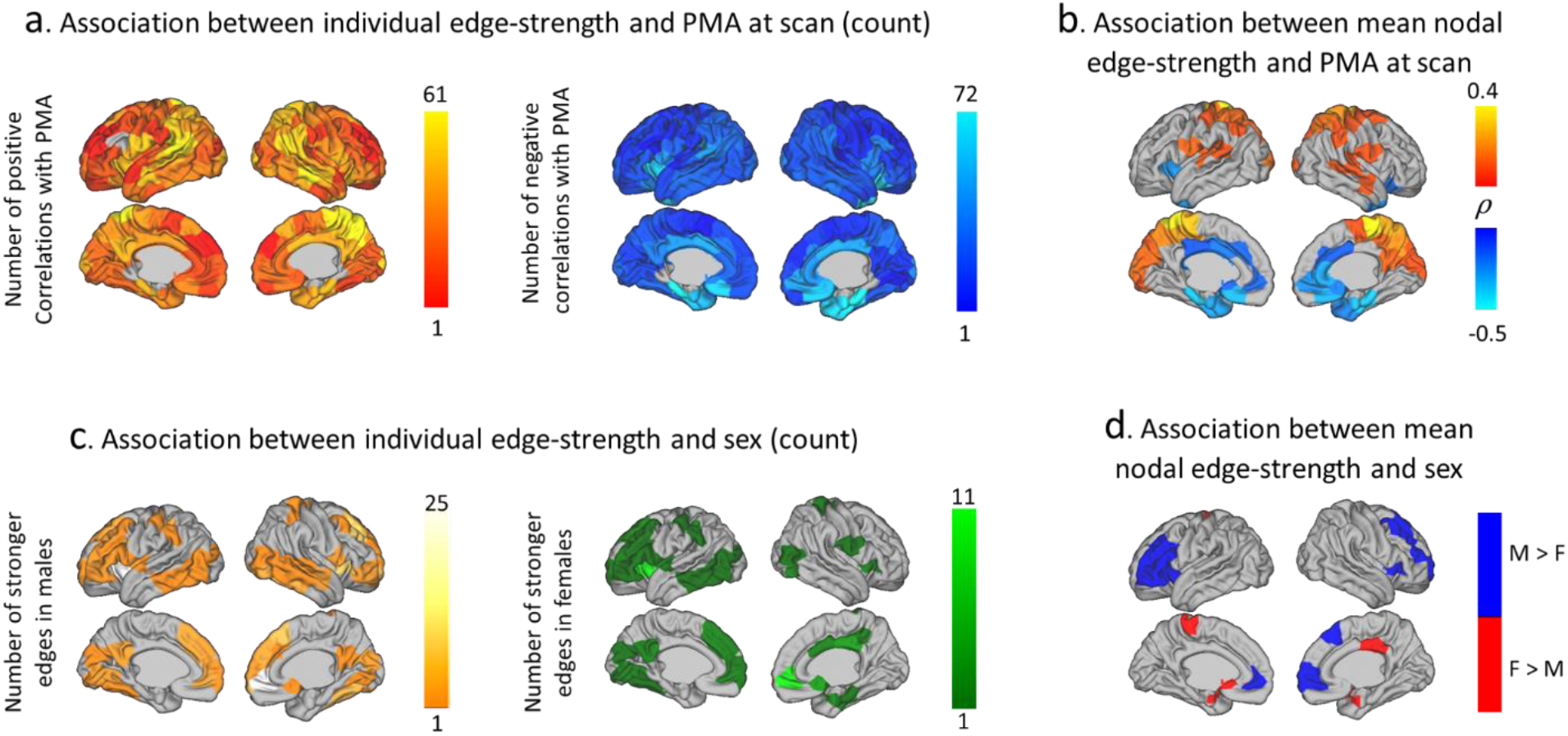
PMA and sex associations with MSNs: a. Sum of positive and negative correlation between inter-node edge-strength and postmenstrual age (PMA) at scan b. Spearman’s correlation between PMA and mean nodal edge-strength c. Sum of stronger inter-node edges in males (orange) and in females (green) d. Significant sex differences between mean nodal edge-strength. All results shown corrected for false discovery rate (FDR) at 5%.

### Modularity of Neonatal MSNs

Using affinity propagation to perform clustering resulted in 12 broadly symmetric cortical modules, aligned with sensory-motor, fronto-temporal, anterior frontal, limbic, cingulate, insular and visual systems (Figure 4 left). Comparable spatial findings were also found when the initial parcellation included different number of regions (n=50,100,150,200,250,300) but the number of clusters increased linearly with parcellation density. For clarity, results are presented only for the middle density (n=150).

**Figure 4.**
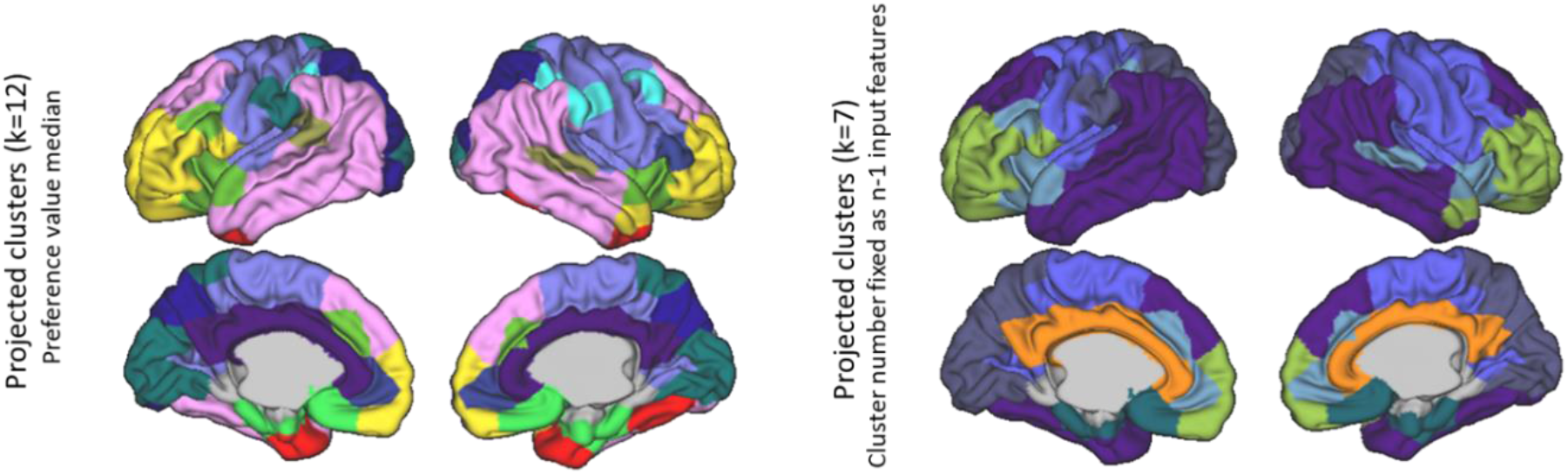
Clustering solution for MSNs created using eight features: Affinity propagation clustering based on a fixed preference value (median similarity) is shown on the left and for a fixed number of clusters on the right.

A similar spatial partition was observed when the number of clusters was fixed at seven (Figure 4 right) and across parcellation densities (see Supplementary Figure 1). To confirm the stability of this solution, clustering was calculated using bootstrapping of 20 random subjects each time, for 500 iterations, and individually for each subject. Node coincidence (how often one node fell in the same node as another) was very high (Supplementary Figure 2b) and even in MSNs of individual infants, the consistency was very good (Supplementary Figure 2a) with the whole sample cluster structure clearly evident in individuals and bootstrap samples.

Comparing the resulting MSNs (for k=7) and von Economo cytoarchitectural classes, similarities were found, more remarkably in limbic and primary sensory areas, as compared to association areas (Figure 5).

**Figure 5:**
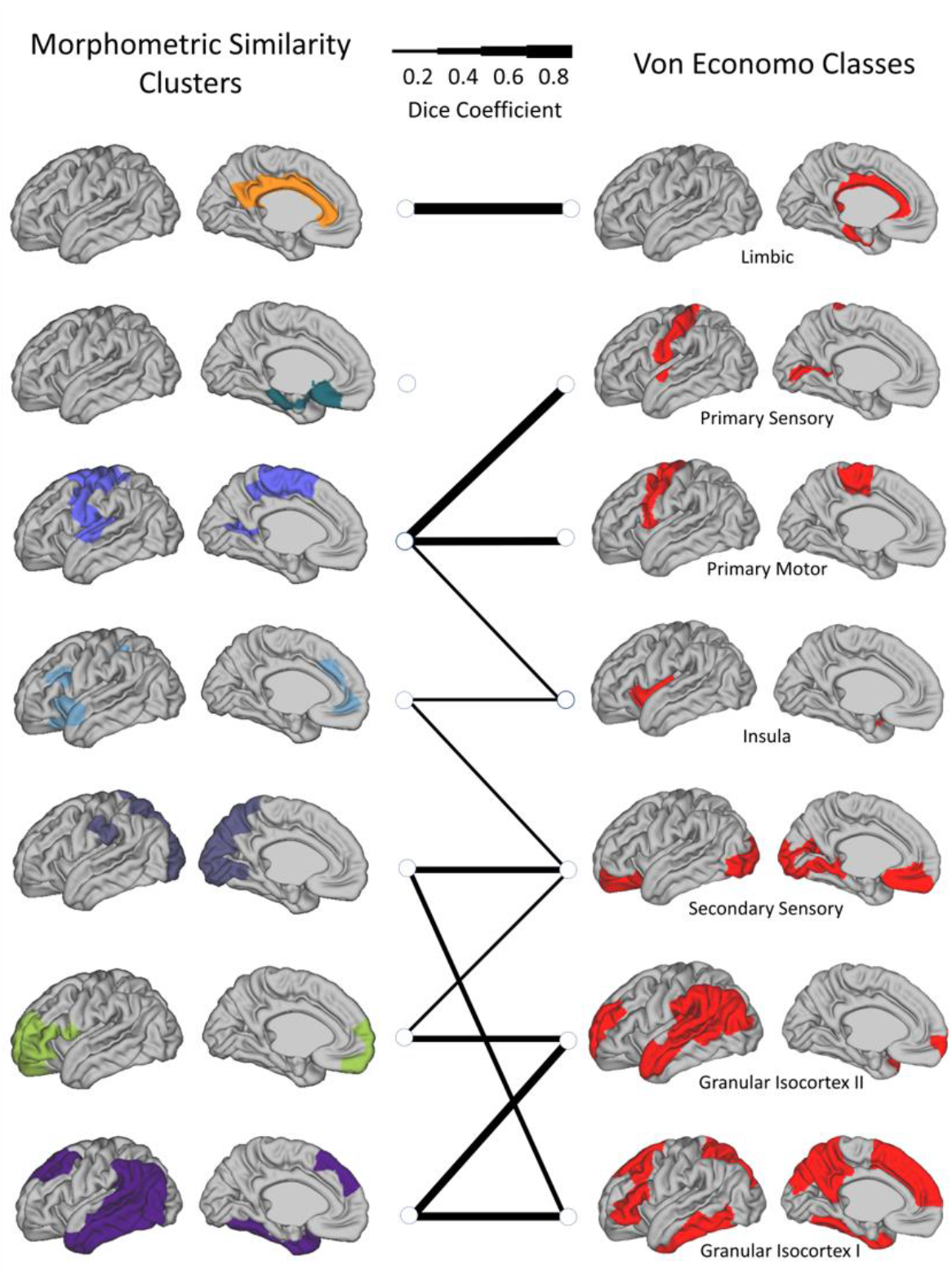
The spatial overlap between MSN modules (left) and von Economo tissue classes (right) quantified as pairwise Dice coefficients. Line thickness indicate the Dice coefficient per pair of connected regions and a Dice of <0.2 is not shown.

#### Feature Contribution

Examination of the clustering solution (for k=7) for each of the *single* morphometric measures, demonstrated that clustering of FA covariance had the highest agreement with the eight-feature solution (normalized variation of information (nvi), z=0.73), while clustering of MC covariance had the lowest agreement (z=0.94) (Figure 6a). The top four metrics in particular are strongly correlated with each other cross-sectionally, derived from the same base diffusion weighted images and therefore this may indicate redundancy. This order remained when the single-feature SCNs were calculated with partial correlation, taking into account PMA, suggesting this is not exclusively derived by age effects in the individual modalities (data not shown).

**Figure 6.**
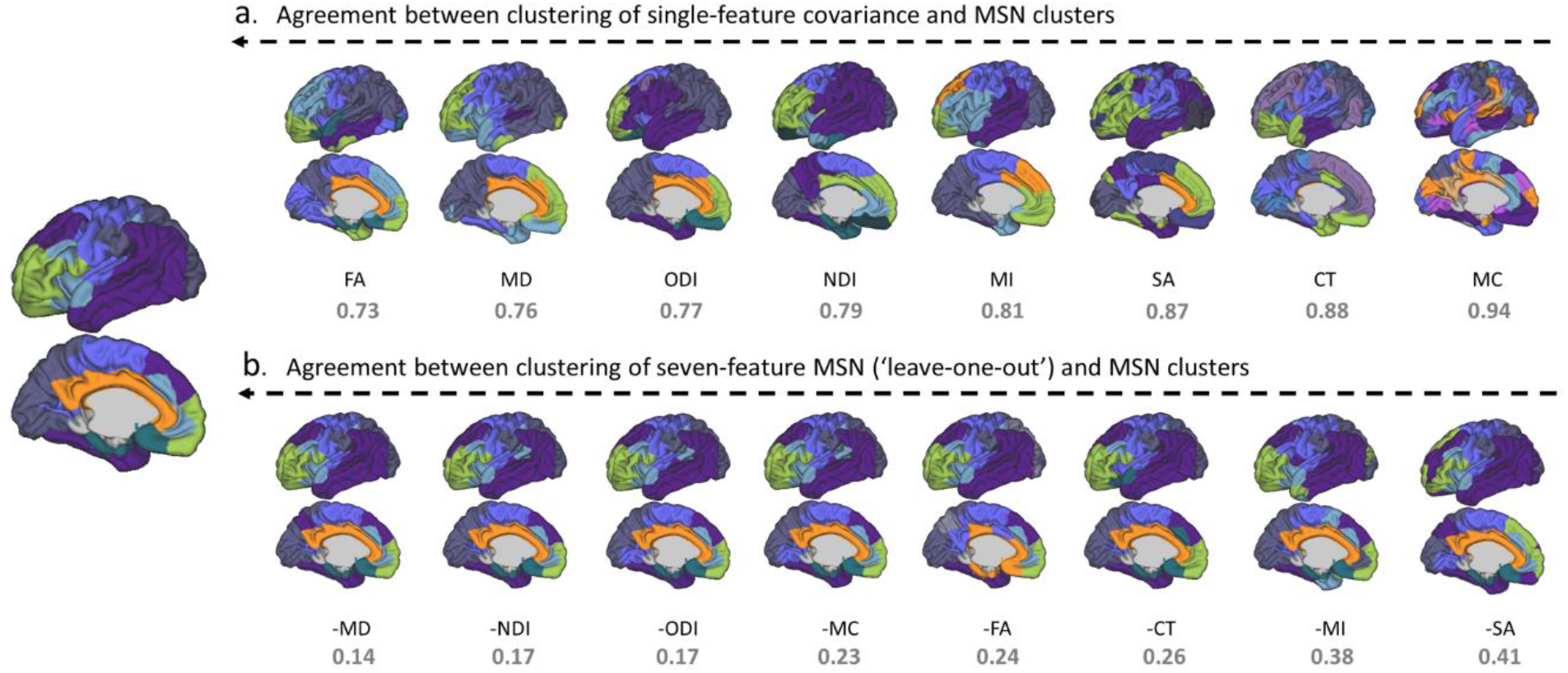
Agreement between clustering solutions using all features (left) with single- (a; top row) and multi-measure (b; bottom row) covariance. Results shown for left hemisphere for ease of interpretation. Clustering solutions are ordered from left (most similar) to right (least similar). **a** Level of agreement between modules derived from a single feature covariance matrix and the eight-feature MSN; **b** Level of agreement between clustering solution for seven-feature MSN (‘leave-one-out analysis’) and eight-feature MSN. In all cases the normalized variation of information value is shown below (lower is better). CT-cortical thickness, MC-mean curvature, MI-myelin index, SA-surface area, FA-fractional anisotropy, MD-mean diffusivity, NDI-neurite density index, ODI-orientation dispersion index.

In the ‘leave-one-out’ seven-feature solutions, the agreement between modules was much higher than for single modalities, as might be expected. Removing SA and MI from MSN estimation had the most profound effect (e.g. resulted in more different clusters) indicating their higher importance to the eventual solution. Removing either MD, NDI or ODI had less of an effect, with near identical solutions (Figure 6b).

#### MSN clusters and PMA at scan

To demonstrate the PMA-related changes in cluster differentiation over time, we performed correlations between individual measures of within-module similarity and between-module similarity. Between-module analysis revealed six cluster-pairs with significant positive correlation with PMA, showing between-cluster integration over time: Occipital-parietal & anterior frontal, occipital-parietal & fronto-temporal, limbic & somatosensory-auditory, limbic & cingulate, anterior-frontal & cingulate and insular-medial frontal & cingulate, while eight cluster-pairs showed significant negative correlation with PMA. The limbic and fronto-temporal clusters demonstrated increased within-module connectivity with PMA, suggesting cluster distinction, or increased internal similarity, while no within-module negative correlation with PMA was found (Figure 7).

**Figure 7.**
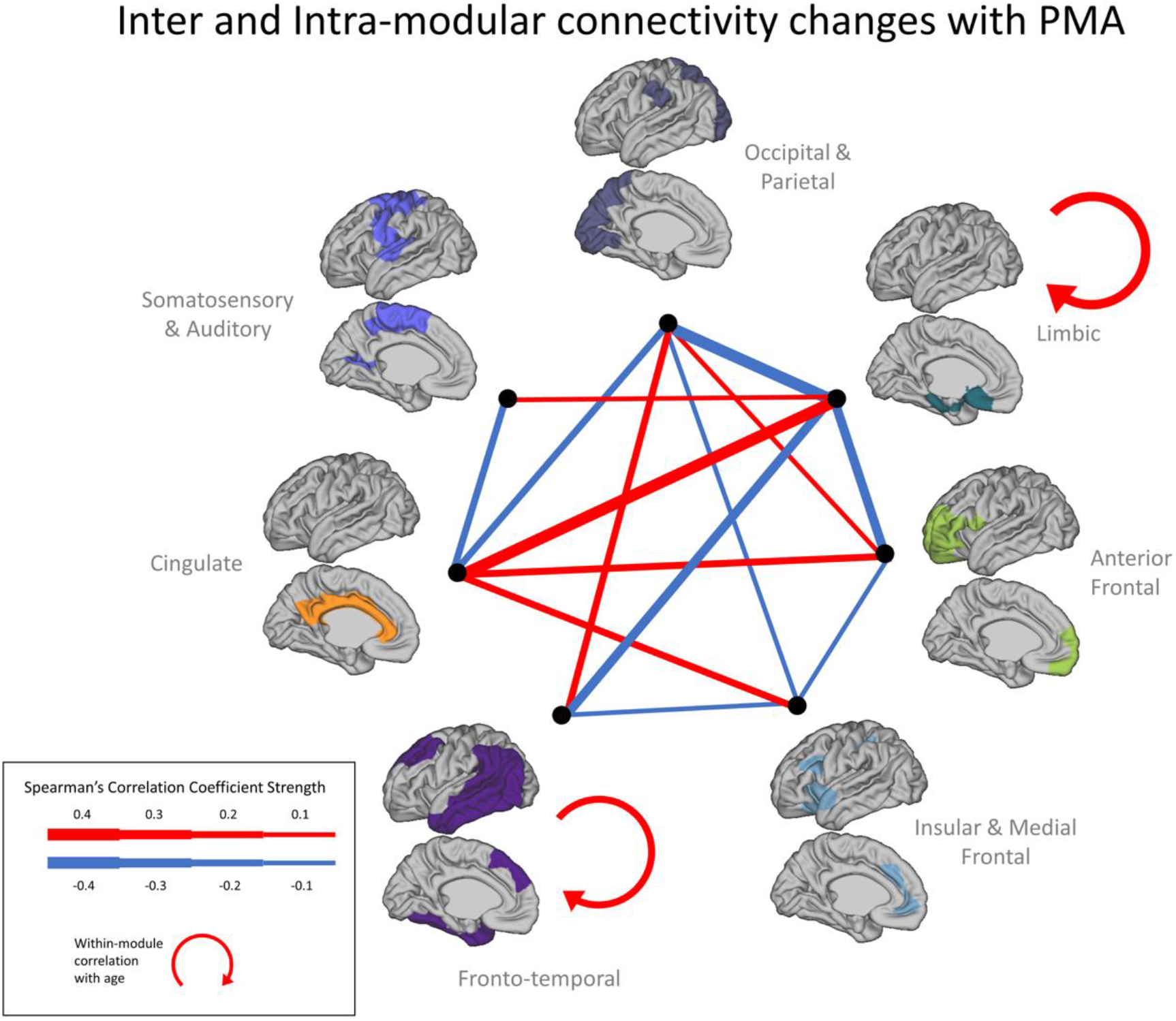
Association between inter-module edge-strength of each pairs of clusters (k=7) and postmenstrual age (PMA) at scan: width of line is indicative of the strength of significant association (results shown corrected for false discovery rate (FDR) at 5%). Correlations between PMA and *intra*-module connectivity is also demonstrated only for the fronto-temporal and limbic clusters.

#### PMA-related associations with clustering solution

Given the clear association between PMA at scan and connection strength between edges and clusters, we investigated the resulting MSN modules when calculated on edge-by-edge age trajectories. We performed clustering on the correlation matrix of each node-to-node MSN edge against PMA. This solution demonstrated both a clearly different result (Supplementary Figure 3), and also has a different interpretation, showing areas that are similar in their direction and strength of maturation. These results showed more long-range connections and much more heterogeneity in frontal cortex. Primary sensory regions clustered together and bilateral symmetric fronto-temporal and fronto-parietal networks were also evident.

## Discussion

Cortical microstructure and morphology go through extensive changes postnatally. Though the majority of neurons are in situ at term age, myelination and synaptogenesis continue to refine the cortical architecture that will persist throughout the lifespan. To profile this architecture in the perinatal brain, we combined multiple structural imaging features, and showed extensive maturational changes throughout a short period of just eight weeks with a posterior-anterior gradient. We demonstrated clear modular network structure, with both local and distributed networks of brain regions having similar cortical profiles. This modular structure showed hemispheric symmetry and mapped onto known cortical functional distinctions.

Diverse refinements in cortical architecture were evident over this short eight-week period of rapid normal development. PMA had a complex effect on edge-strength showing both positive and negative correlations, i.e. regional profiles became more and less similar with PMA. These age-related associations were confined to occipital, parietal and temporal areas and were less evident in later maturing frontal areas. Whereas these neocortical associations were positive (more connectivity with the rest of the brain), limbic and anterior cingulate regions showed negative correlations with PMA (less connectivity). These findings suggest that immediately following birth, sensory and limbic areas and posterior parietal regions as opposed to regions related to higher executive functions (e.g. prefrontal cortex) have the largest maturational changes, qualitatively similar to the pattern seen in Lebenberg et al. (2019) with older infants. Changes in these executive regions may be evident in higher-order functional areas only at a later stage, in line with the more prolonged synaptogenesis, dendritic arborization and myelination patterns (Huttenlocher, 1990; Huttenlocher & Dabholkar, 1997; Teffer & Semendeferi, 2012). This may also reflect work in cortical GM volume maturation in neonates, showing a posterior-anterior gradient in the first weeks following birth (Gilmore et al. 2007).

In a small number of infants using MSNs based on structural and extended diffusion metrics, but not surface measures, Galdi et al. (2019) also found that brain structures became both more similar and dissimilar to each other over the same PMA range. While that study used different feature vectors, atlas and populations to here, some findings are clearly complementary: Occipital connections were found to increase with PMA, parietal edges were also more positively than negatively associated with PMA, while temporal edges showed mixed associations with PMA. In contrast to our findings, frontal regions also showed a strong relationship with PMA, however this was more evident in white matter (WM) rather than GM.

When examining each morphometric measure in isolation, developmental trajectories with clear anatomical structure were found across the PMA range examined replicating prior work (Batalle et al., 2019; Dean et al., 2017). Typical measure of cortical anatomy, thickness and area, as well as a good measure of relative T1w contrast, myelin content, showed strong increases with age (Li et al. 2013). Age however showed little significant association with curvature, in agreement with previous reports showing relatively little change in MC compared to other surface measures in the first months following birth (Batalle et al. 2019; Li et al. 2015), suggesting that cortical folding remains relatively static in this period. Though no single individual measure mirrored the clustering solution achieved by combining all measures, the four diffusion metrics had the better correspondence (Figure 6a), but were apparently less important to the clustering solution (Figure 6b). This apparent contradiction is due to redundancy. For example, MD and NDI show very similar patterns of correlation with PMA (Figure 2), so removal of one or two diffusion measures does not reduce the effective information used to calculate correlations in MSNs.

Cortical diffusion metrics show complex maturational changes before and after the period considered as term birth. Ball et al. (2013) and Batalle et al. (2019) have shown that MD decreases linearly with PMA, while FA decreases rapidly until week 38 PMA, but then begins to slowly increase. This might explain why at the PMA range examined here (37-44 weeks) we observed both negative and positive associations between FA and PMA. Moreover, Batalle et al. found that NDI is positively associated with PMA, similar to the results here. ODI seemed to stabilize after 38 weeks, whereas we still observe a strong association with PMA after 38 weeks. We might hypothesize that this is related to the focus on preterm neonates in that work, where microstructural trajectories could differ due to earlier birth and the very different early environmental exposures that entails (both clinical and ex-utero related).

Sex differences were less widespread compared to age effects. While sex differences in cortical volumes are evident from birth (Gilmore et al. 2007), they are subtle when looking at single modality covariance networks in early development (Geng et al., 2017; Zielinski et al., 2010). Using the MSNs approach, we were able to detect sex effects in distributed cortical areas, most pronounced in frontal and temporal regions. This reflects similar regions in another neonatal study using tensor based morphometry (Knickmeyer et al., 2014) and even in adults, multimodal imaging sex differences (combination of T1w, T2w and diffusion-weighted images) were mainly localized in the frontal lobe, followed by parietal and temporal lobes (Feis et al., 2013); So this may represent a consistent gross pattern throughout the lifespan.

Generally, any network construction and follow-up clustering analyses are highly reliant on several factors, including population and age range examined (Morgan et al., 2019; Seidlitz et al., 2018), type of morphometric measures used (Nie, Li, & Shen, 2013), parcellation (Arslan et al., 2018), threshold (Bordier, Nicolini, & Bifone, 2017), wiring cost (Betzel et al., 2017) and type of modularity estimation (Sporns & Betzel, 2016), and are therefore likely to vary accordingly. For module definition, we applied affinity propagation to discover meaningful cortical clusters, without eliminating or changing any connections for network construction. In young adults, utilizing MSNs and the Louvain algorithm, Seidlitz et al. (2018) reported four cortical modules consistent with lobular division. Using affinity propagation to cluster similar developing regions in terms of CT and curvedness between the ages 3-20, Nie, Li, and Shen (2013), found eight and five clusters, respectively, also with some alignment to lobular division, and indeed our parcellation based just on CT was qualitatively similar.

While most of the resulting modules presented in these studies tend to be spatially contiguous and local, our clustering solution showed also long-range connections, and might be more indicative of possible functional and structural connectivity. The modules were broadly aligned with known functional systems, with relative stability when fixing number of clusters to seven or allowing the algorithm to select itself (k=12). These included sensory-motor, fronto-parietal, temporal, limbic, cingulate and visual regions. While functional brain networks are evident at this early stage, higher-order adult-like systems will only fully emerge at a later age (Gao, et al., 2015; Keunen, Counsell, & Benders, 2017). Our correspondence between MSNs and functional systems may reflect the underlying architecture for later developed functional networks (Geng et al., 2017; van den Heuvel et al., 2015). Moreover, our resulting MSN clusters show some overlap to the von Economo tissue classifications, implying of the possible origins of the cortical profiles obtained in our analysis and provide some reassurance of their validity. Further supporting this, the solution revealed here also shows several analogies to the clustering solution of genetic contribution to SA and CT reported by Chen et al. (2013) and might be especially relevant in early development where external environmental effects on these parameters are still relatively small.

The clustering solutions were spatially robust, by and large not affected by PMA (confirmed by results of clustering separately only individuals scanned in the top and bottom PMA-range quartiles), nor was it affected by the neonates’ ex-utero experience, as the clustering solution for neonates scanned within a week from birth resembled that of the entire sample (data not shown). So although PMA did not alter the location of structure of the MSNs, it was associated with their internal coherence. PMA was linked to increased connectivity within the clusters and both increased and decreased connectivity between clusters (cluster integration and segregation). Though the exact pattern was complex, the summary was that cingulate showed integration with the limbic and insular clusters while segregating from most of the neocortical clusters, perhaps suggesting an advancement towards a more broad representation of paralimbic structures, while the limbic and fronto-temporal clusters showed also PMA-dependent increase in intra-similarity. Increases in network segregation in the developing brain has been previously demonstrated in neonates (Cao et al., 2017; Zhao et al., 2019) and in children and adolescents (Baum et al., 2017; He et al., 2018), and may show more distributed integrative patterns in adulthood (Fair et al., 2009).

While our study’s advantages include a large term-born neonatal sample with data acquired and analyzed with optimized protocols for this age group, as well as the use of a large number of morphometric measures to characterize the brain, it is not without limitations. In order to describe developmental growth patterns, longitudinal, instead of cross-sectional data is needed. Due to the nature of the morphometric measures utilized in this work, we were not able to assess whole brain connectivity; here excluding subcortical structures, cerebellum and brain stem. Future work should investigate the structural and functional coupling of neonatal MSNs, as well as their genetic and environmental correlates. The implication of recent work seeing changes in MSNs in common and rare neurodevelopmental disorders (Morgan et al., 2019; Seidlitz et al., 2019) is that these changes must occur very early in development. Using such a large sample, our study provides a means to detect meaningful changes and alterations in structural coupling at an early stage in infants with either a known genetic or high likelihood to have a later neurodevelopmental disorder.

To conclude, in this study we describe cortical MSNs in the neonatal brain, their association with PMA and their community structure. We report that following birth, PMA is strongly related to edge-strength, in both a positive and negative manner, and that these associations show a posterior-anterior gradient. The contribution of individual morphometric measures, as well as their developmental trajectories, are variable, supporting their combination in order to more accurately estimate brain maturation and connectivity. Clustering of MSNs was related to known functional systems and was found to be relatively stable during the PMA range examined, while within-cluster strength generally seemed to increase postnatally, implying increased network segregation in this short period.

## Acknowledgements

The developing Human Connectome Project was funded by the European Research Council under the European Union Seventh Framework Programme (FP/20072013), Grant Agreement no. 319456. Infrastructure support was provided by the National Institute for Health Research (NIHR) Mental Health Biomedical Research Centre at South London, Maudsley NHS Foundation Trust, King’s College London and the NIHR Mental Health Biomedical Research Centre at Guys, and St Thomas’ Hospitals NHS Foundation Trust. The views expressed are those of the author(s) and not necessarily those of the NHS, the NIHR or the Department of Health and Social Care. The study was supported in part by the Wellcome Engineering and Physical Sciences Research Council Centre for Medical Engineering at King’s College London (grant WT 203148/Z/16/Z) and the Medical Research Council (UK) (grants MR/K006355/1 and MR/LO11530/1). D.F’s PhD is supported by the Sackler Institute for Translational Neurodevelopment at King’s College London and the Medical Research Council Centre for Neurodevelopmental Disorders, King’s College London. J.O.M. is supported by a Sir Henry Dale Fellowship jointly funded by the Wellcome Trust and the Royal Society (Grant Number 206675/Z/17/Z). J.O.M., D.E. and G.M. received support from the Medical Research Council Centre for Neurodevelopmental Disorders, King’s College London (grant MR/N026063/1).

