## Supplementary Material for "Development of Microstructural and Morphological Cortical Profiles in the Neonatal Brain"

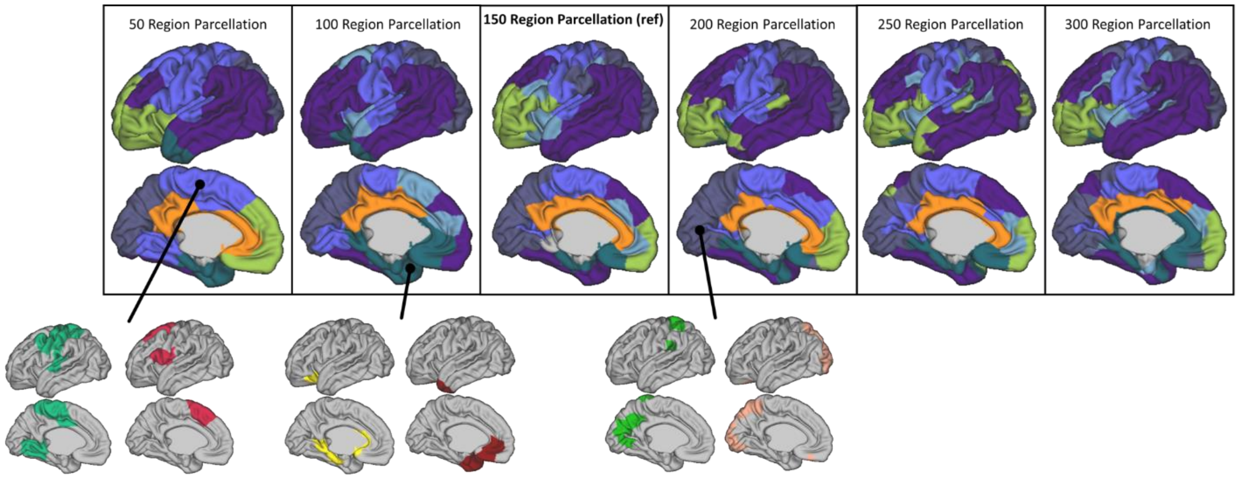


**Supplementary Figure 1:** Clustering with a fixed k=7 across different parcellation densities. Colours are matched to the best overlap parcel in the n=150 parcellation. In some cases two clusters match maximally to only one cluster in the n=150 clustering solution. These are indicated in the bottom row.

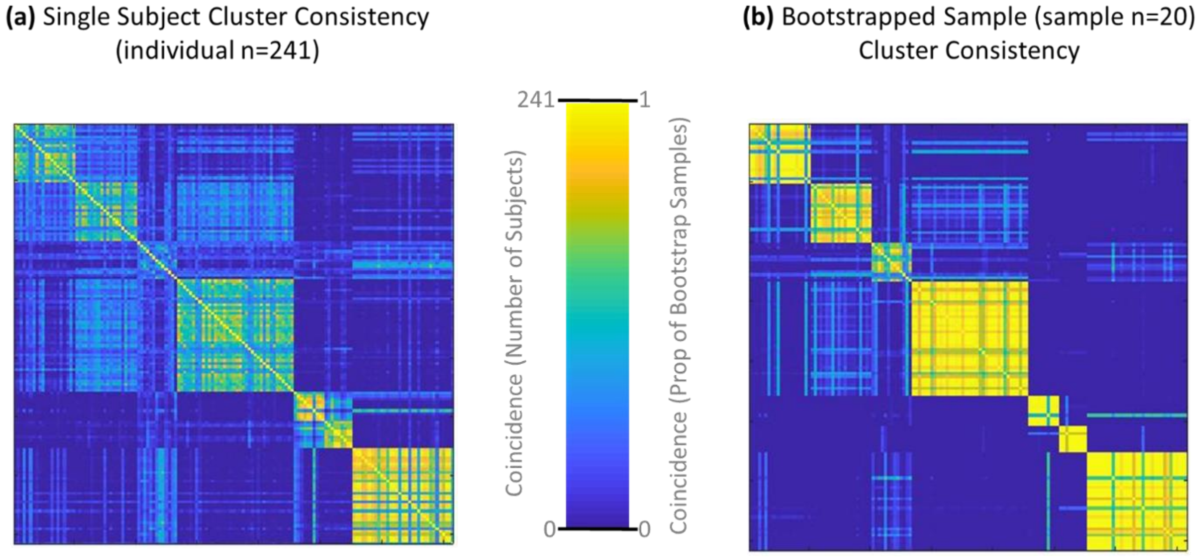


**Supplementary Figure 2:** Cluster repetition with a fixed (k=7) clustering repeated for each individual (a) and for 500 bootstrap resamples of 20 subjects (b). Each element in the matrix indicates the count (a) or proportion (b) of times each node (row) was co-incident in a cluster solution with each other node (column).

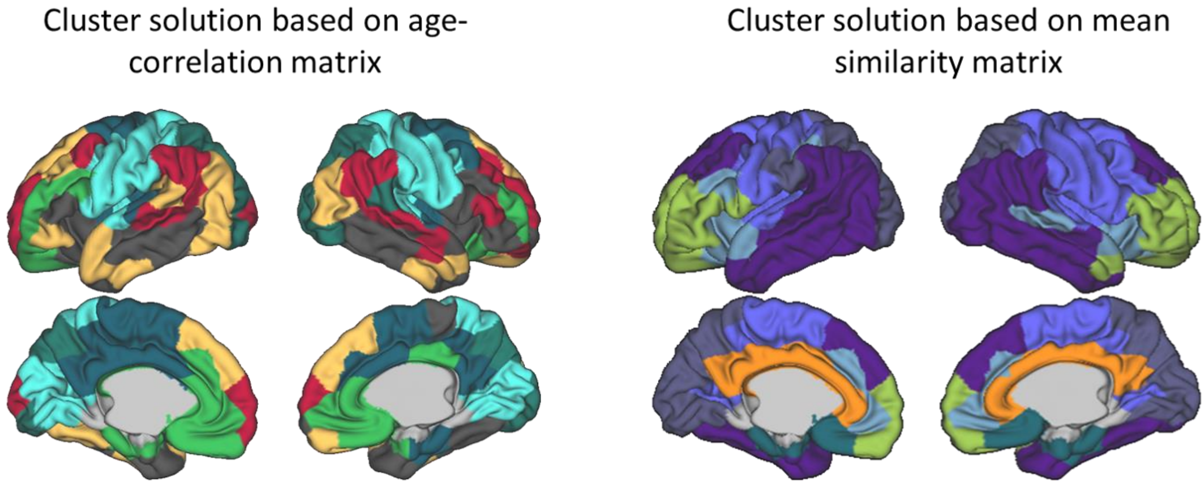


**Supplementary Figure 3:** Comparison of clusters derived from the mean MSN across subjects (a) and the correlation matrix of the MSN edges against PMA (b). Both have k fixed to 7.
